# Super-enhancer interactomes from single cells link clustering and transcription

**DOI:** 10.1101/2024.05.08.593251

**Authors:** Derek J. Le, Antonina Hafner, Sadhana Gaddam, Kevin C. Wang, Alistair N. Boettiger

**Author notes:** These authors contributed equally (Derek J. Le, Antonina Hafner).

## Abstract

Regulation of gene expression hinges on the interplay between enhancers and promoters, traditionally explored through pairwise analyses. Recent advancements in mapping genome folding, like GAM, SPRITE, and multi-contact Hi-C, have uncovered multi-way interactions among super-enhancers (SEs), spanning megabases, yet have not measured their frequency in single cells or the relationship between clustering and transcription. To close this gap, here we used multiplexed imaging to map the 3D positions of 376 SEs across thousands of mammalian nuclei. Notably, our single-cell images reveal that while SE-SE contacts are rare, SEs often form looser associations we termed “communities”. These communities, averaging 4-5 SEs, assemble cooperatively under the combined effects of genomic tethers, Pol2 clustering, and nuclear compartmentalization. Larger communities are associated with more frequent and larger transcriptional bursts. Our work provides insights about the SE interactome in single cells that challenge existing hypotheses on SE clustering in the context of transcriptional regulation.

## Introduction

Enhancers play a pivotal role in transcriptional control in higher eukaryotes, binding cell type-specific transcription factors and activating distal cognate promoters to provide the context-specific gene expression needed for development and for responses to changes in environmental signals ^1^. Enhancer-mediated gene activation depends on some degree of proximity between the enhancer and target promoter, which is constrained by the 3D organization of the genome. Traditional models suggest that such regulatory interactions generally occur in a pairwise manner, in which individual enhancers contact individual genes ^2,3^. However, recent studies have suggested that some enhancer-based regulation involves multi-enhancer contacts, including interactions from extremely distal enhancers (>10 Mb) and *trans* contacts (for review ^4–9)^. This hub model challenges the conventional understanding of enhancers as largely pairwise and largely locally regulated.

Interest in hub-mediated regulation has spurred the development of new technologies to detect multi-way interactions. For example, Tri-C ^10^, TM3C ^11^, COLA ^12^, C-Walks ^13^, MC-4C ^14^, Pore-C ^15^, triplet-Hi-C ^16^, GAM ^17^, and SPRITE ^18^ have all reported preferential three-way interactions among enhancers. Among these clustered enhancers, the large, highly occupied enhancers, sometimes termed super-enhancers ^19,20^ (SEs), are most enriched ^10,14–18^. The term SE was first introduced in mouse embryonic stem cells (mESCs) to define a superlative group among enhancers with disproportionately high levels of transcription factors (OCT4, SOX2, and NANOG) and Mediator ^19^ in concentrated genomic windows of a few to a few tens of kilobases, which tended to be near genes with important roles in cell identity ^19,20^.

However, it is currently unclear at what absolute frequencies and physical distances enhancer clustering occurs as existing methods only measure enrichment relative to other types of multiway interactions. If enhancer clustering occurs in only 1% of cells, it is unlikely that the clustering plays a major role in regulating transcriptional bursts, regardless of how “enriched” the clustering is. Differences in length scale have substantial implications for distinguishing clustering and regulatory mechanisms, as highlighted in recent debates and reviews ^4–9,21^. However, it is unclear from these sequencing assays at what length scale the reported clustering occurs, and how different the length-scales of proximity assayed in these methods are. Sequencing-based approaches to measuring clustering must also contend with recovery bias and amplification bias ^22^. Technical parameters, such as crosslinking time, fragment length, or slice size, further constrain the analysis of elements in a cluster, with statistical power additionally confined by the number of cells and the recovery rate of the multi-way interactions^23^.

It is also unclear from recent work how (and if) multi-way enhancer clustering affects gene expression. This gap has spurred development of new multi-modal single cell-omics technologies, including methods that can find multi-way contacts in single cells, such as scSPRITE ^24^, and a new scHi-C ^25^; measure pairwise DNA contacts along with RNA expression, like HiRES ^26^ and GAGE-Seq ^27^; and even measure multi-way contacts and RNA in the same cells, like R&D SPRITE ^28^, MUSIC ^29^, DNA-seqFISH+^30,31^ and DNA MERFISH ^32^. However, these emerging methods currently lack the sensitivity and coverage to explore associations between enhancer hubs and transcription bursts. Concurrently, protein clusters have been seen to associate with transcriptional bursts in fixed ^20^ and live cells ^33,34^, though it is unclear if these clusters involve enhancer hubs or individual loci. Recent reviews have especially highlighted the important gaps left in recent evidence of transcription hubs in testing what impact clustering has on transcription of associated genes ^4–9^.

To overcome these constraints, and quantify enhancer clustering we employed genome-scale DNA fluorescent *in situ* hybridization (FISH) using the optical reconstruction of chromatin architecture (ORCA) technique ^35^. We focused on a comprehensive set of previously defined “super-enhancers”, as this group includes many of the enhancers we find most interesting and is as superlative in its clustering ^10,14–18^ as in its TF occupancy ^19,20^ and its roles in cell identity ^19,36–39^, suggesting SEs are a good model of enhancer clustering. We mapped the SE interactome from mouse embryonic stem cells and related its organization to the transcription of key pluripotency genes. Our interrogations revealed that, while SE contacts are rare–i.e. most SEs are isolated from one another in most cells–SEs do assemble at larger length scales we call ‘communities.’ These communities assemble cooperatively under the combined effects of genomic tethers, TF clustering, and preferential localization to nuclear speckles. Interestingly, larger communities are associated with more frequent and larger transcriptional bursts, linking community-level, not contact-level, clustering with gene expression.

## Results

### Global Spatial Organization of SEs in Single Cells

To elucidate the spatial characteristics of SE clustering, we conducted a comprehensive mapping of the 3D spatial positions of all SEs (**Figure 1A**). We combined all SEs from the seminal work on SEs in mouse embryonic stem cells (mESCs) by Whyte et al., with an additional 145 elements elevated as SEs in a later classification by Sabari et al., resulting in a master list of 376 SEs (**Table S1**). We note that several of the previously defined SEs span TSSs (see **Table S1**) and thus both enhancers and some promoters are included in our assay. To precisely locate the positions of these SEs at the single-cell level, we targeted each SE with a unique sequence tag by *in situ* hybridization and detected these tags with independent fluorescent oligos in independent images using ORCA ^35^. This approach is more laborious than DNA-MERFISH and related methods ^30,32^, which assay many targets per image, but it allows spatially clustered signals to be more reliably detected and identified. Probes were designed by selecting 15 kb regions centered on the previously identified SE calls (**Figure 1B**, **Table S1**), ensuring uniformity in our experimental design and detection efficiency.

**Figure 1.**
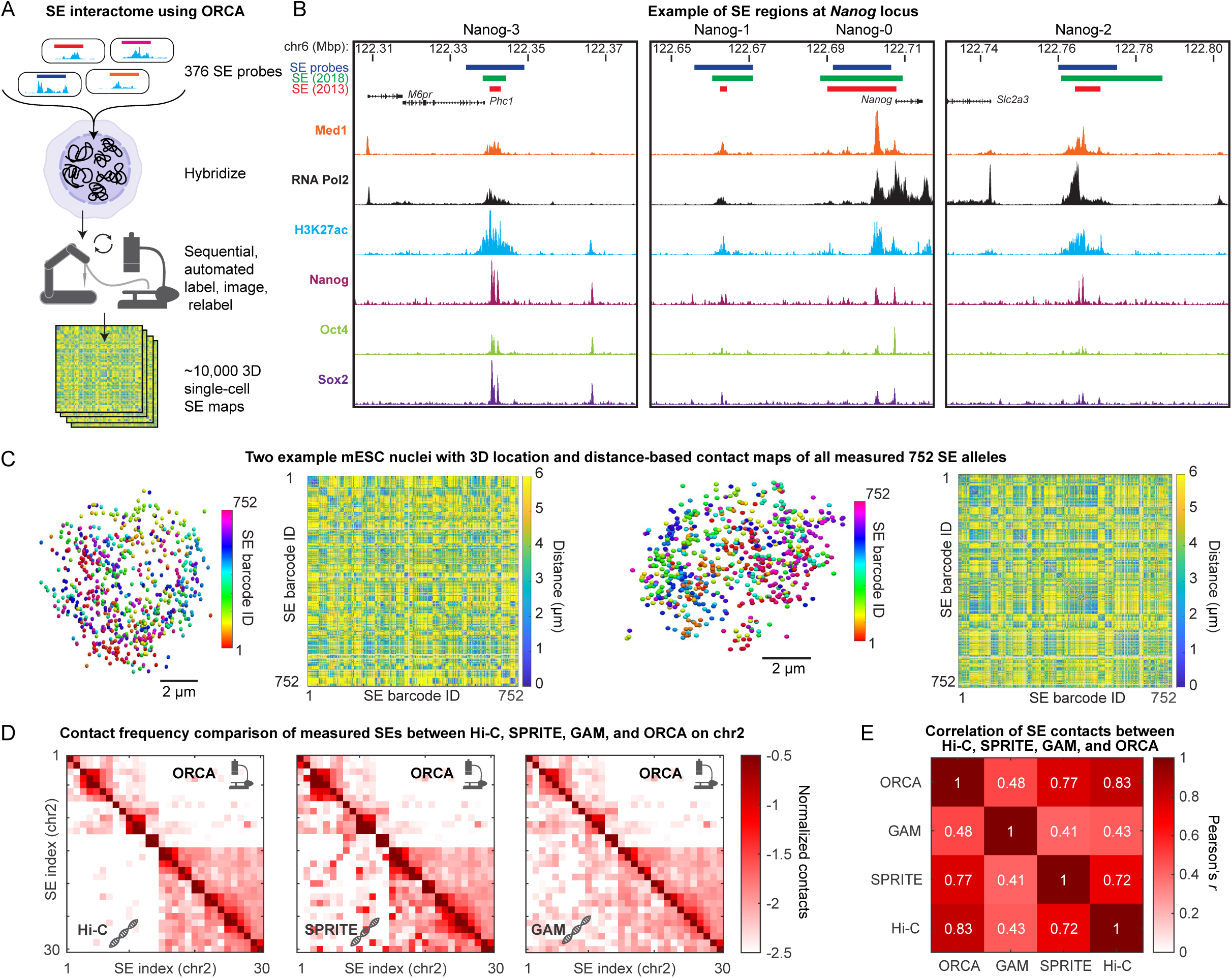
ORCA enables visualization of all mESC SEs across single cells. **A.** Schematic of approach, where each SE is individually labeled and imaged. Two SEs can be labeled with distinct fluorophores and imaged in one round with 188 rounds to image all 376 SEs. **B.** Genome browser tracks of *Nanog*-associated SEs. SE annotations by Whyte et al. (2013) and Sabari et al. (2018) are shown in red and green respectively. 15 kb regions, centered on Whyte annotations, were picked for ORCA and are shown as blue boxes. ChIP-seq of Med1, RNA Pol2 (Sabari et al. 2018), H3K27ac (Sethi et al. 2020), Nanog, Oct4, and Sox2 (Whyte et al. 2013) of transcription factors and histone modifications. **C.** Examples of two cells showing the positions of the two alleles of the 376 imaged SEs (752 SEs total per cell) and the corresponding single cell SE-SE pairwise distance maps. **D**. Comparison of relative contact frequencies among SEs on chromosome 2 (all other chromosomes shown in Figure S2) measured by ORCA (<300 nm threshold), Hi-C (Bonev et al. 2017), SPRITE (Quinodoz et al. 2018) and GAM (Beagrie et al. 2017). **E**. Correlation of contact frequencies for *cis* contacts across all chromosomes between ORCA, Hi-C, SPRITE and GAM.

We imaged more than 5,000 cells across in over 400 sequential imaging rounds (including control images used for quantifying measurement uncertainty and correcting for chromatic aberration). These probes allowed us to discern 752 distinct loci per cell, accounting for both alleles of each SE (**Figure 1C**). Following corrections for technical errors including stage drift, sample drift, and chromatic aberrations, we achieved high spatial precision, with a residual error of ∼60 nm (**Figures S1A-S1D**). During image processing, 14 SEs were dropped from the analysis due to limited detection efficiency or additional off-target binding, while the remaining SEs exhibited largely uniform and highly efficient detection (median >90%) (**Figure S1E**). Overall, our comprehensive imaging experiment measured over 2 billion inter-SE distances across 343,000 images and 6 TB of data, providing a rich foundation for further exploration into the intricate landscape of SE-SE interactions.

To validate our ORCA dataset, we computed pairwise SE interactions maps for each chromosome (in *cis*) and conducted a thorough comparison between ORCA and three previously employed genome-wide assays: Hi-C ^40^, GAM ^17^, and SPRITE ^18^. Hi-C involves *in situ* ligation of restriction-digested genomes to identify 3D proximity among sequence elements. GAM utilizes whole-genome sequencing of isolated nuclear cross-sections to reveal patterns of sequence clustering, avoiding the need for restriction digestion or proximity ligation. SPRITE employs a split-pool barcoding strategy to identify crosslinked sequences within the same nuclear fragment, also avoiding the use of proximity ligation. All three methods offer a relative measure of interaction frequencies across the entire SE-interactome examined in our data, while the differences between the methods offer a measure of control to account for method-specific artifacts or biases. Our findings reveal a robust, highly significant correlation between the SE ORCA data and all three comparison methods (**Figure 1D**). Notably, all three methods exhibit a stronger correlation with ORCA than with each other (**Figure 1E**). Assuming that shared features among the four techniques reflect the true biology of mESC SE interactions and that divergent features indicate technique-specific artifacts, this comparison suggests that ORCA exhibits the least artifacts, underscoring its reliability in capturing mESC SE interactions.

### Super-enhancer contacts are rare and involve few SEs

To determine when SEs are likely in contact, it is instructive to know the physical size of a typical SE, and thus understand what center-to-center distances between SEs are consistent with contact. The well-studied Sox2 Control Region (SCR) SE has been previously imaged with probes tiling the ∼15 kb SE region in 5 kb steps ^35,41^. We found that the median distance between the two adjacent probes to be ∼175 nm, while probes at opposite ends of the single SE were ∼225 nm apart, in our measurements and in independent results from Huang et al ^41^ (**Figure 2A**). Based on these measurements, we decided to use 200 nm as a cutoff distance for calling SE contacts.

**Figure 2.**
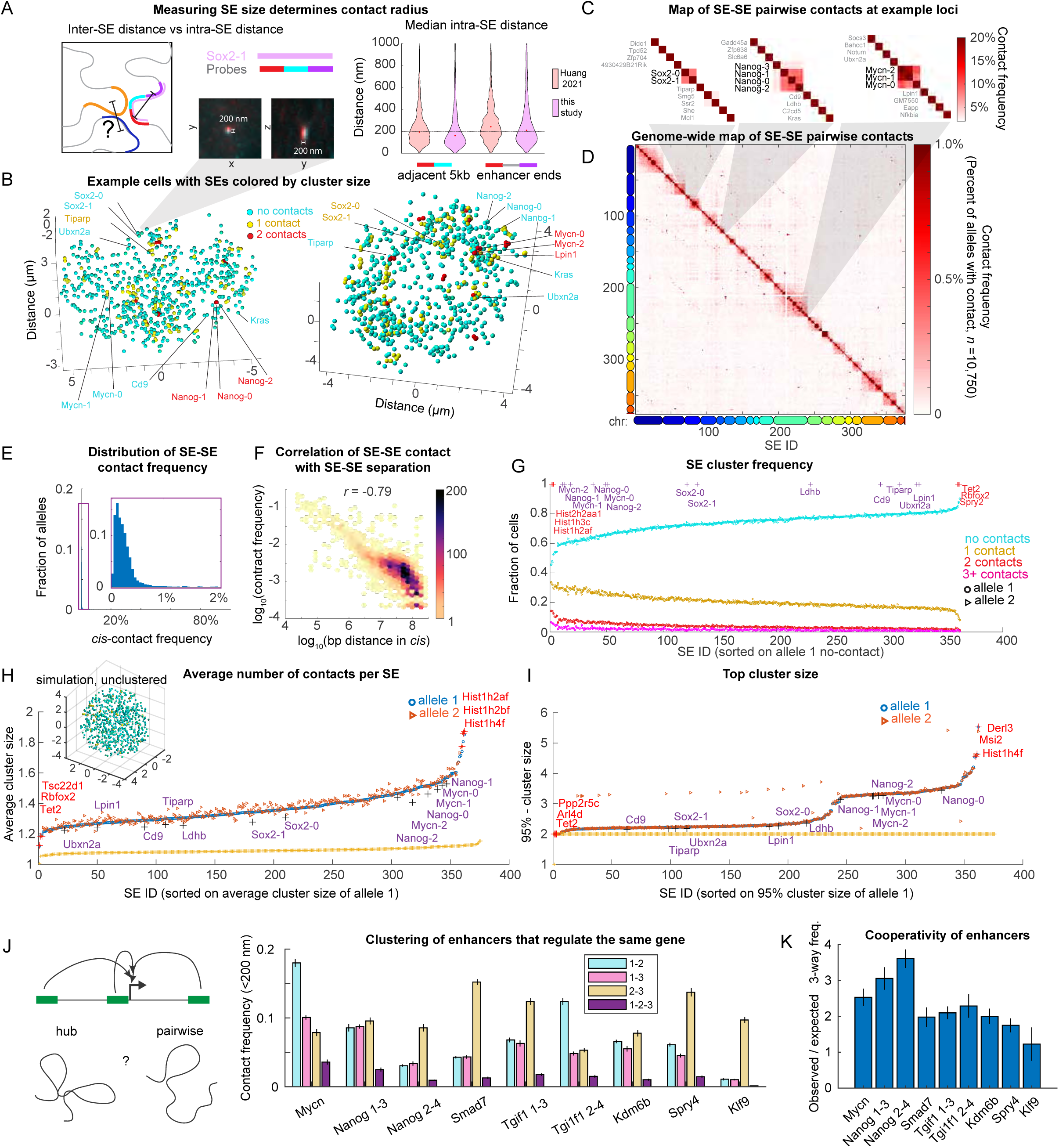
Multiway SE contacts are infrequent. **A.** Schematic illustrating the approach of determining the distance threshold for SE-SE contact. Colored bars show 15 kb SE probes and the Sox2 SE tiled with 3 adjacent 5 kb probes. Quantification of the interprobe distance of adjacent and end-to-end pairs by Huang et al. and this study (*n*=430, *n*=2,359, *n*=479, *n*=1,312). **B**. Examples of two single cells with positions of all imaged SEs colored by the cluster size (number of other SEs within 200 nm). **C**. ORCA contact frequency (percent over all alleles) between pairs of SEs around *Sox2*, *Nanog* and *Mycn*. **D**. Pairwise map of all SE contacts (<200 nm), with positions of SEs in C. highlighted. **E**. Distribution of contact frequency for all *cis* SE-SE pairs. **F**. Correlation between genomic separation between SEs (in *cis*) and contact frequency. Heatmap represents the density of points. **G.** Fraction of cells exhibiting 0, 1, 2, 3, 4+ SE partners, across all cells. SEs are sorted by the fraction of cells with 0 contacts. **H.** Average contact-cluster size for each SE. SEs are sorted by average number of contacts of the first allele. Average contact-cluster size was also calculated for simulated cells with 752 randomly positioned SEs (orange line) and the example of a simulated cell is shown in the inset. **I**. Number of SEs in the 95-percentile largest contact-cluster it forms, sorted from smallest to largest. The 3 highest and 3 lowest SEs per analysis are marked with a red ‘+’ and named, and the positions of example SEs around *Sox2*, *Nanog* and *Mycn* SEs are marked with a purple ‘+’ and named. **J.** Two-way and three-way contact frequency among SE triplets that regulate the same target gene. **K**. Cooperativity in 3-way contacts among SE triplets that regulate the same target gene. Error bars represent +/- one standard deviation, estimated by bootstrapping with *n*=5,375 cells.

Examining individual SE alleles from individual cells, we were surprised to find few pairwise SE-SE contacts and even fewer multi-way contacts (**Figure 2B**). Nonetheless, SE-SE interactions of known regulatory importance did exhibit more frequent contact. For example, both the *Sox2* promoter region and its 115 kb distal *SCR* are SEs in our SE interactome, and deletion of the *SCR* or reduction of this interaction leads to reduced *Sox2* expression ^41–43^. This SE pair exhibited contacts in 9% of the alleles analyzed. Similarly, the promoter-linked SE of *Mycn* and its more distal SE also exhibited contacts in 8% of alleles, while the *Nanog* promoter-linked SE (*Nanog-0*) and its distal SEs (*Nanog-1* and *Nanog-2*) ^24,44^ were in contact in 10% of alleles (**Figures 2B and 2C**). In contrast, across all *cis*-linked SEs (SEs on the same chromosome) in all cells, contacts between any particular pair were rare, with the vast majority occurring in less than 1% of alleles (**Figures 2D-2F**).

Despite this low contact frequency with specific pairwise partners, we hypothesized that with hundreds of SEs to choose from across the genome, many SEs may still be in contact with some other SE much of the time. To test this hypothesis, we computed the overall frequency for each SE to form pairs, triplets, or clusters of four-or-more with any other SE (**Figure 2G**). We found all SEs, save a handful associated with histone genes, exhibited no discernible contacts with any other SEs in 60-80% of cells. In 20-30% of cells, these SEs were in contact with one other SE (**Figure 2G**), greatly surpassing the frequency of specific pairwise *cis-*interactions (median 0.2%, **Figure 2E**) and indicating a considerable degree of variability in SE partnering. SEs formed multiway contacts of 3 or more SEs in 5-10% of cells and formed contacts of 4 or more SEs in <5% of cells (**Figure 2G**). To further quantify cluster size we also computed the average cluster size per SE (**Figure 2H**) and the 95th-percentile largest cluster formed by each SE across all cells (**Figure 2I**). This confirmed that all the SEs were alone on average (cluster size 1.2 - 1.6) and rarely formed hubs (95th-percentile largest cluster being only 2-5 SEs).

This rare degree of contact was nonetheless higher than expected based on simulations of nuclei with random positioning of 752 SEs (representing two alleles for each measured SE) in the same nuclear volume (**Figure 2H**, inset), where the average cluster size is ∼1 (**Figures 2H and 2I**).

Thus, SEs exhibit a non-random degree of clustering (consistent with prior claims ^15–18,28^), but are on average isolated from each other and not in SE contact hubs. That contact hubs are so rare does not exclude the possibility that they have an important function, but it does exclude a model in which the dominant mode of contact occurs through hubs.

While most SEs looked similar in terms of their average number of contacts with other SEs, our deep dataset allowed us to detect subtle yet significant differences (**Figures 2G-2I**). The highest average contacts were exhibited by SEs near the replication-dependent histone genes, a known enhancer cluster that was also detected by SPRITE ^18^. SEs with known functional interactions (e.g. *Sox2* SEs, *Nanog* SEs, *Mycn* SEs, see **Figure 2C**), ranked among the highest interacting SEs, whereas other SEs (e.g. *Cd9* SE, *Ldhb* SE, the next most proximal SEs to *Nanog*) did not (**Figures 2C and 2H**).

Some genes are regulated by three-or-more super-enhancers in the same cell type (**Figure 2J**). In our SE-interactome, twenty-three SEs are part of regulatory triplets, targeting 7 unique genes. We asked if these SEs formed gene-specific interaction hubs. From these triplets, 3-way contact hubs were observed in 1-3.5% of cells, which was 2-10 fold less often than pairwise contacts (**Figure 2J**). For example, *Nanog*-0, *Nanog*-1 and *Nanog*-2 were found in a 3-way hub in 3.5% of cells, though pairwise contacts between the promoter linked *Nanog-0* and the distal *Nanog*-1 or *Nanog*-2 occurred in 10% of cells. This indicates that when multiple SEs regulate the same gene, their contacts largely occur at different times, rather than through a single hub. Nonetheless, for all the genes regulated by 3 or more SEs, the three-way contact was observed at a higher frequency than expected if contacts had been statistically independent, evidencing some cooperativity (**Figure 2K**). In the example provided by the *Nanog* SEs, the 3-way interaction would be expected in only 1% of cells if it were truly independent.

The tendency of SEs to cluster more than other genomic elements has been repeatedly found by sequencing assays, and here our single-cell imaging approach has allowed the frequency to be placed on an absolute scale. Furthermore, this analysis demonstrates that, somewhat unexpectedly, hub formation (contact cluster) is not the typical state for an ESC SE. Given that most of the genes associated with SEs are highly expressed (transcribed in more than 5% of cells) much of that transcription activity must occur when the SE is not in a contact cluster. Even SEs that regulate the same target do not form constitutive hubs. Nonetheless, an infrequent (<5%) but significant fraction of the time, most SEs can be found in contact clusters (**Figure 2I**), allowing the potential that clustering provides transient contributions to enhancer function.

### Super-enhancers form communities

SEs (and cellular components in general) may not necessarily need direct physical contact in order to influence each other’s function ^45–48^ ^49–51^, raising the possibility that SEs may have spatial organization beyond the 200 nm scale relevant for their behavior. Despite the rarity of contacts, visual inspection of SE organization in our data suggests a degree of clustering at larger scales not captured with our contact-analysis (based on the 200 nm threshold) (**Figure 3A** contrast to **Figure 2B**). We termed this SE clustering without requiring contact “SE communities”. We quantified the frequency and specificity of community formation across our dataset using an arbitrary 600 nm threshold (see **Figure S3** for alternate thresholds). This larger threshold is comparable to the size of many nuclear condensates (e.g. Med1 and Pol2) ^20,52,53^, and various nuclear bodies such as speckles ^54^.

**Figure 3.**
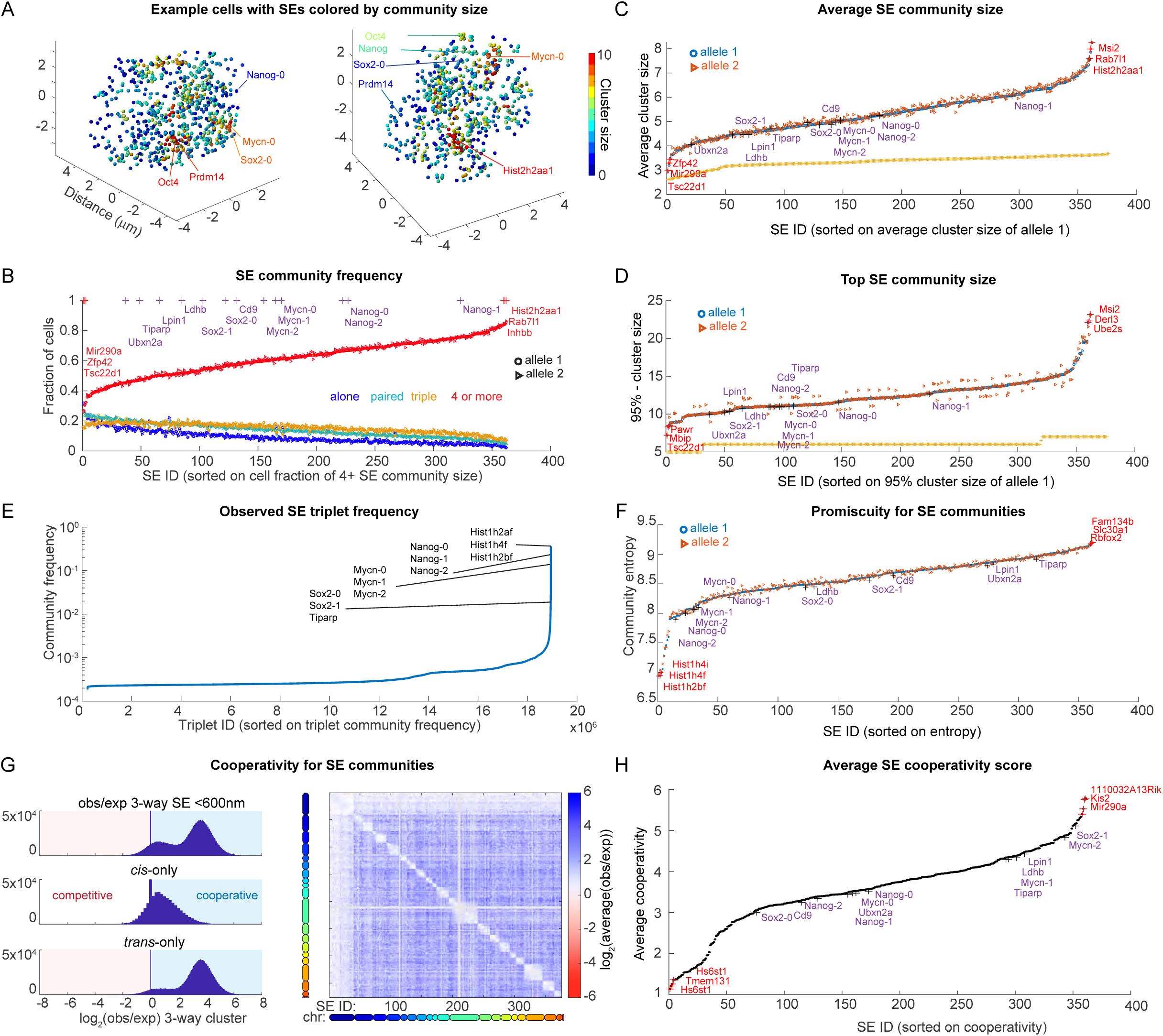
Presence of SE communities in the cellular landscape. **A.** Visualization of two single cells with the positions of all SEs color-coded by community size (number of SEs within 600 nm). **B**. SE cluster frequency (600 nm) for each SE. **C.** Average community size sorted by the first allele (‘**o’**). Second allele (‘D’) for simulated cells with 752 randomly positioned SEs. **D**. 95th percentile largest SE community size for each SE, with SEs ordered by their 95th percentile community size. The 95th percentile largest community size of simulated cells with randomly positioned SEs is shown in orange. **E**. Frequencies of all detected three-way contacts within 600 nm. Triplets are sorted by their detection frequency. **F.** Computation of entropy for each SE over all community interactions across cells as a measure of promiscuity. SEs are ranked by their entropy score, with higher entropy indicating greater promiscuity. **G.** Cooperativity (log_2_(O/E)) for all 3-way interactions (left). Map of average cooperativity among SE triplets in community formation. For each pair, we computed the observed versus expected cooperativity with a third SE. Heatmap shows average over all possible 3rd SE. **H.** Average cooperativity per SE (averaged over all possible values of the 2nd two partners). The 3 highest and 3 lowest SEs per analysis are marked with a red ‘+’ and named, and the positions of example SEs around *Sox2*, *Nanog* and *Mycn* SEs are marked with a purple ‘+’ and named.

At the community scale, and in contrast to our observations at the contact scale (**Figure 2G**), we found that the majority of SEs were not alone (**Figure 3B**), with nearly all SEs found in communities of at least 4 SEs in over 40% of cells. The SE community size was significantly larger than expected for a random distribution of SEs (**Figure 3C**). For most SEs their top 5% largest community included up to 10-14 SEs, with a few SEs found to be in communities of up to 20 (**Figure 3D**).

To assess the diversity of community members associated with each SE, we initially examined the frequency of specific triplets. Over two-thirds of all observed triplet combinations (approximately 20 million) were unique, occurring only once in 10,000 cells. Less than 10% of all triplets were observed more than 10 times in 10,000 alleles (**Figure 3E**). Nonetheless, a handful of triplet combinations were more common, with specific triplets of different histone enhancers occurring in over 30% of cells. Communities containing triplets of SEs in recognized regulatory interactions (3 *Nanog* associated SEs*, 3 Mycn* associated SEs) were also observed at relatively high frequency (1-10%) compared to other SE triplets (<0.1%) (**Figure 3E**). Given that nearly all SEs had an average community size of >4, the rarity of specific triplets indicates a low degree of specificity within the communities.

To further quantify the specificity vs randomness of SE community composition, we computed the entropy for each SE across the population of cells (**Figure 3F**). An entropy value of zero indicates that an SE interacts with a specific set of SEs in its community in every cell. Conversely, an entropy value of ∼9.6 (which is log_2_(752)) indicates a random choice (uniformly distributed). The relatively high entropy values for most SEs (ranging from 8 to 9) imply SE community associations are promiscuous. However, histone-associated SE clusters deviated from this trend, displaying lower promiscuity (entropy values of ∼7) compared to the majority of SEs (**Figure 3F**). Additionally, entropy exhibited a modest anti-correlation with average community size (Pearson’s *r*=-0.52), indicating that the most isolated SEs primarily form communities through chance encounters, while the least isolated ones maintained a more stable core community (**Figure S3A**). Thus community membership is generally diverse and promiscuous, though the size and variability of composition vary measurably among SEs.

We next asked if community assembly is cooperative. To quantify cooperativity, we compared the frequency of the observed 3-way association to the expected frequency if association was not cooperative (i.e. association of the third element is independent of whether the first two associate). Globally, we found most SE triplets formed cooperatively (**Figure 3G**), on average ∼8x more than expected (95th percentile = 19x more), though certain triplets showed little cooperation (5th percentile =1.000x). Averaging across the third triplet member allowed us to map these interactions (**Figure 3G**, right), revealing that *trans* interactions were generally more cooperative than *cis*. This is likely a polymer effect: while any two given chromosome territories are rarely in contact, once an SE is close to one SE from another chromosome it is more likely to form a community with other SEs also from that chromosome. Nonetheless, even *cis* interactions showed a strong cooperativity (**Figure 3G**). We then compared per SE its average cooperativity with all other SEs. The *Sox2* and *Mycn* promoter-proximal SEs were especially cooperative, as was *Mir290a* - a strong super-enhancer that typically formed among the smallest communities, and *Hist1h2an*, a super-enhancer from the histone complex which typically forms among the largest communities (**Figure 3H**).

For thresholds of ∼400-700 nm we observed similar patterns to those described with the 600 nm threshold. All SEs on average form a community, the community size depends on SE identity, and the ranking of community size correlates with the ranking found for 600 nm (**Figures S3B and S3C**).

In summary, we found that at the community scale (clustering without necessarily being close enough for physical contact), SEs are not spatially isolated, in contrast to our observations at the contact scale. Furthermore, we noted substantial SE-specific differences in terms of their community size, specific membership, and cooperativity of assembly.

### Factors influencing community size

We next investigated correlates of community size to understand potential factors influencing community formation and explaining SE-specific differences in average community size (**Figure S4**). The genomic tethers between SEs may increase the probability of encounters and contribute to larger communities among genomically clustered SEs (**Figure 4A.i**). Transcription factors enriched at SEs, including SOX2, POL2, and Mediator, have large IDRs that self-associate into liquid condensates ^20,52,53,55,56^ and are hypothesized to cluster enhancers without nanoscale contacts ^4–9^ thus, they may contribute to community formation (**Figure 4A.ii**). The chromatin polymer has been hypothesized to undergo a polymer-phase separation based on histone modification state in which H3K27ac regions cluster, providing another potential mechanism to assemble SEs (**Figure 4A.iii**). Prior work has also shown that SEs tend to avoid LADs ^57^, and tend to associate with nuclear speckles ^18,24,28,58,59^, raising the possibility that preferential migration to or from a common nuclear region may drive community formation (**Figure 4A.iv**).

**Figure 4.**
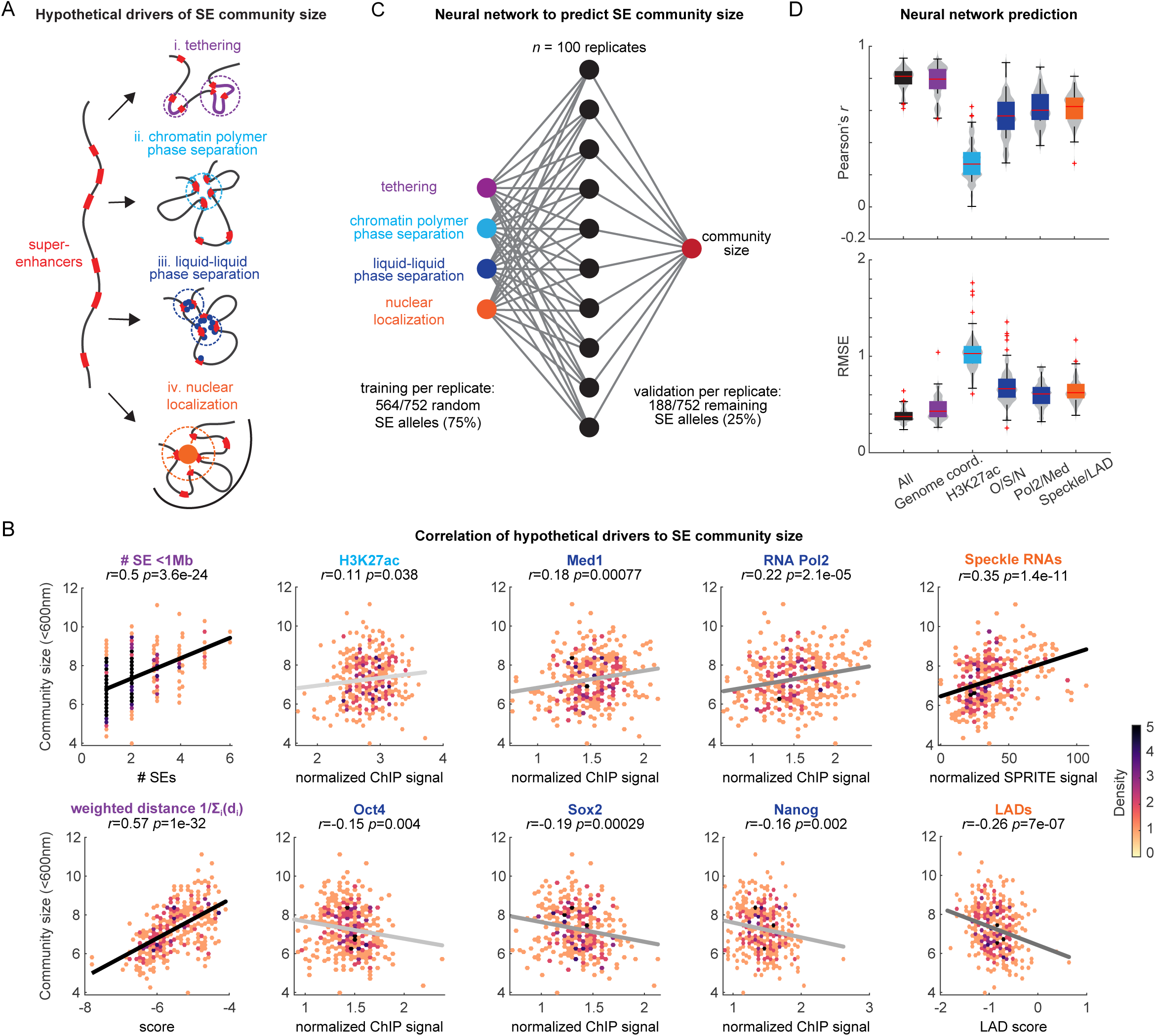
Identification of SE characteristics that determine community size. **A.** Schematic illustrating potential mechanisms facilitating community formation (i) chromatin tethers between genomically proximal SEs, (ii) chromatin heteropolymer phase separation into homotypic domains, (iii) liquid-liquid phase separation of proteins that bind enhancers, and (iv) migration to a common nuclear landmark. **B**. Pearson correlation across SE features in (A), and the average community size for each SE. Gray/black lines are linear fits to the data, darker line colors reflect smaller p-values from the Pearson correlation. **C**. Schematic representation of the neural network (NN) used to integrate all the SE features in (A) and (B) to predict the average SE community size. **D**. Top: Pearson correlation coefficient between the NN predicted community size and the measured community size calculated for models run with either all SE features as the input or just individual or a subset of features (indicated on the x-axis). Bottom: Root mean square error between the NN prediction and the measured community size. O/S/N: *Oct4*/*Sox2*/*Nanog*.

Both the number of SEs within 1 Mb and the weighted sum of the inverse distance between all SEs correlated with community size (*r*=0.51, *r*=0.56), indicating that the linear organization of the genome plays a substantial role in community formation but is not the only factor (**Figure 4B**). By comparison, H3K27ac ChIP-seq levels showed no significant correlation with community size (**Figure 4B**), while levels of SOX2, OCT4 (POU5F1), or NANOG ^19^ all showed weak negative correlation with community size (**Figure 4B**), arguing against a K27ac/polymer-driven or TF-induced clustering. However, ChIP-seq levels for components of the general transcription machinery, POL2 and MED1, both showed significant positive correlation (**Figure 4B**) (*r*=0.22 *p*=3e-5, *r*=0.16, *p*=0.0025), consistent with the recent hypothesis that POL2 and/or MED1 condensates may contribute to enhancer clusters without inducing direct contact ^20,52,53^.

To explore the contribution of sub-nuclear localization tendency to community formation, we used publicly available lamin DAM-ID from mESCs ^57^ as a measure of the frequency of lamin association for each of the SEs and publicly available RNA-DNA SPRITE data from mESC with speckle-specific RNA factors ^28^ as a measure of association with speckles. Community size showed significant anti-correlation with LAD association frequency (*r*=-0.24, *p*=5.2e-6) and positive correlation with speckles (*r*=0.3, *p*=2.5e-10) (**Figure 4B**), suggesting sub-nuclear positioning also contributes to community formation.

To further investigate how informative all the individual SE characteristics are for its community size, without assuming a linear combination between correlates, we used a machine learning approach. Using a single-layer, fully-connected, neural network (NN) with 10 nodes, and the aforementioned correlates as input variables (**Figure 4C**), we trained the model to predict community size for each SE. The NN achieved an average Pearson’s correlation of *r*_NN_=0.80, with an average root mean square error (RMSE) of 0.38 SEs per community (**Figure 4D**), with averages taken across replicate training rounds.

To understand the relative contributions of different correlates, we then trained identical networks using only a subset of predictive variables. Surprisingly, predictions trained on Pol2 and Med1 alone with no information about SE distances or chromosome identity exhibited a notable degree of accuracy (*r*_NN_=0.62, RMSE = 0.60). Training on combined levels of Oct4, Sox2, and Nanog or combined positional information of Speckle and LAD association also showed improved predictivity, while linear separation still provided the best predictions as expected from our correlation analysis (**Figure 4D**). Taken together, we concluded that SE community size is primarily determined by the linear arrangement of SEs within the genome, with non-trivial, additional contributions arising from binding of transcription-associated TFs such as MED1 and POL2, and sub-nuclear positioning.

### Super-enhancer communities boost transcription

We next investigated if community size affected transcription bursting for SE-linked promoters. We combined ORCA labeling for 44 SEs with RNA FISH for nascent transcription from 6 associated (and highly studied) genes (*Nanog, Sox2, Myc, Pou5f1, Prdm14, Tbx3*). We noticed, in some cells, the allele with a larger SE community exhibited transcriptional bursts (**Figure 5A**).

**Figure 5.**
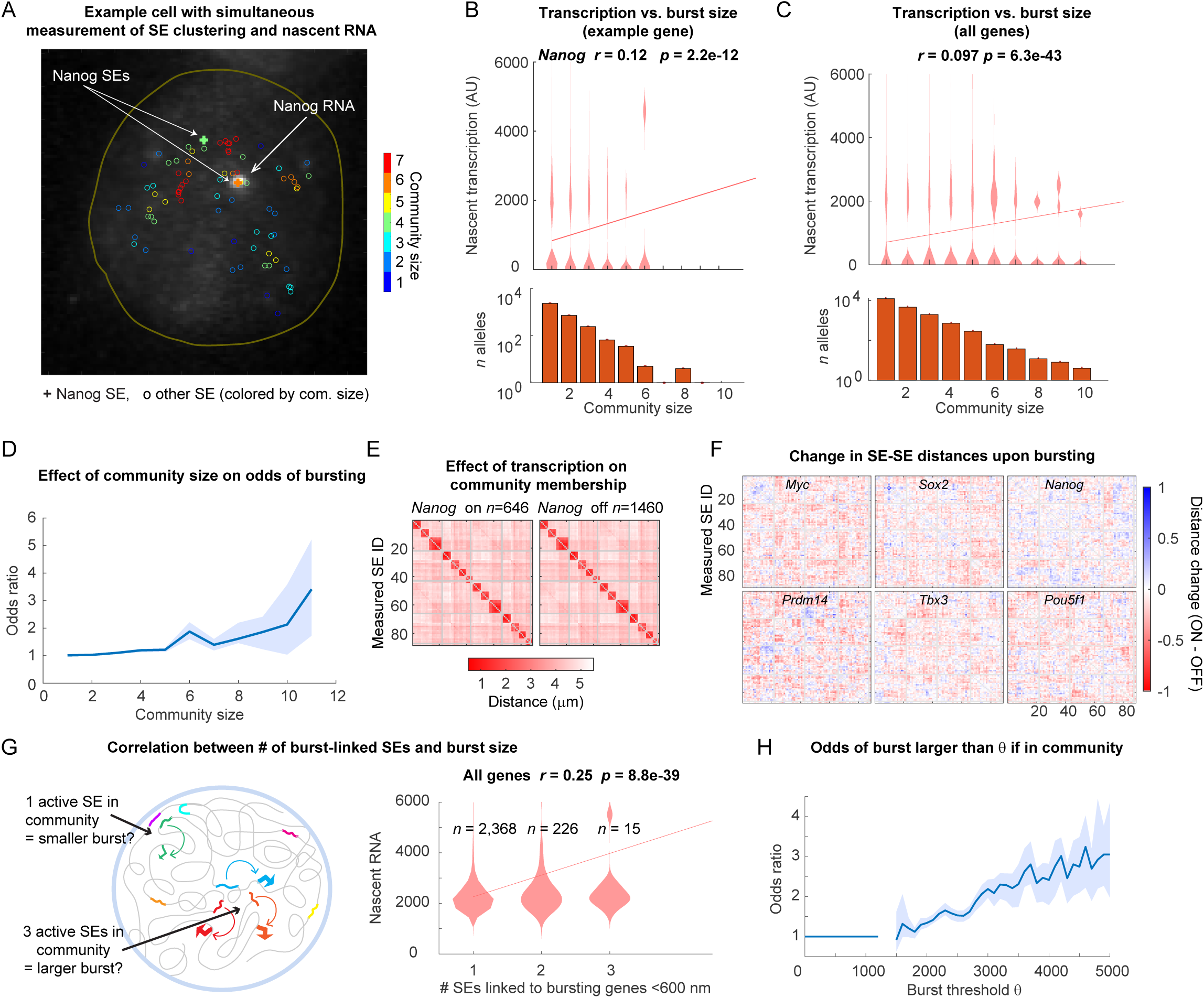
Functional role of SE communities for transcription. **A.** Example of a single cell showing nascent transcription of Nanog (‘Nanog RNA’) together with z-projected positions of SEs. SE are colored by the community size. ‘+’ indicates the position of the SE located at the *Nanog* promoter. **B**. Top: distributions of nascent transcription intensity in arbitrary units (AU) of *Nanog* for different SE community sizes. Pearson’s correlation coefficient and p-values are indicated at the top of the plot and the line over the violins is the best linear regression through all the data. Bottom: number of alleles (number of cells x 2 chromosomes in each cell) observed for each SE community size. Note, top and bottom plots share aligned x-axes. **C**. Same as (B) but aggregating data for the 6 measured genes: *Nanog*, *Sox2*, *Myc*, *Pou5f1*, *Prdm14*, and *Tbx3*. **D**. Odds ratio of active transcription (y-axis) given the indicated community size (x-axis) that the gene-proximal SE participates in. **E**. Median pairwise distance maps between measured SEs in this experiment for cells that are actively transcribing *Nanog* (‘*Nanog* on’) and cells with no *Nanog* nascent transcription (‘*Nanog* off’). **F**. Median pairwise distance difference maps between actively and non-actively transcribing cells for the 6 measured genes in (C). **G**. Left: Schematic of how active SE clustering affects transcription. Active SEs are shown connected to an associated gene (promoter arrow). Right: Violin plots of nascent transcription levels for different numbers of co-transcribing genes within 600 nm. Line over the violin plots shows the linear regression through all points. **H**. Odds ratio of a transcriptional burst being higher than *theta*, when in a community with other transcribing genes.

To quantify these observations, in every cell we measured the nascent transcript intensity, scoring intensity as zero if the gene lacked a detectable punctum within 1 um of the associated SE. We then computed the community size of the associated SE and observed a significant correlation with transcription **(Figures 5B and S5,** *p=*2.2e-22). A similar correlation held when data from all labeled genes was combined (**Figure 5C**, *p*=6.3e-43). To further quantify the effect size of this correlation, we computed the odds of bursting as a function of community size. The Odds Ratio of active transcription for clustered SEs (compared to isolated ones) increased modestly from 1.25-fold higher in communities of size three, to 3-fold higher for the rare communities of size ten (**Figure 5D**), further suggesting that community formation augments transcription. Despite this observed boosting effect, the data also indicate community formation is not sufficient for bursting, as a substantial number of SEs in communities of 3+ members were silent (**Figures 5B and 5C**).

Next, we explored whether SE communities with bursting genes contained distinct subsets of SEs compared to those associated with silent genes. We therefore mapped the median all-to-all distances among SEs for all assayed genes as a function of burst state (**Figures 5E and S5**). These maps looked similar, suggesting transcription dependent differences in global 3D SE organization are minor. However, upon subtracting these maps between transcriptionally on and off alleles, the heatmaps are largely a uniform pink-red, indicating (**Figure 5F**), the bursting population exhibited generally closer SE-SE distances for all SEs for all genes, with little SE-specific pattern (**Figure 5F**). This means, for example, *Pou5f1* is closer to *Sox2* when *Nanog* is bursting (rather than just being closer to *Nanog*), and that the same pattern holds for most possible triplets. This general pattern can be explained by the combination of three things: (1) correlation of clustering with bursting (**Figures 5A-5D**), (2) the multipoint nature of communities, such that when a given SE is in a community, not only is it closer to other SEs, but these other SEs in that community are closer to one-another (3) the fact community identity associated with bursts is a non-specific effect among SEs, rather than something that requires a particular pair or triplet to interact.

Finally, we asked if “active SEs” (here defined as those associated with active transcription bursts) preferentially clustered with other active SEs, and if this had any impact on the average burst size (**Figure 5G**). For each SE associated with active transcription (defined above), we computed the number of other SEs also associated with active transcription within 600 nm. We observed a clear positive correlation between the number of active SEs in a community and the burst size (*r*=0.25, *p*=8.8e-39) (**Figure 5G**). We then asked if the odds for a gene to have a burst size larger than a threshold, *theta*, increased if another gene within 600 nm had a burst size larger than *theta* (**Figure 5H**). We see for *theta* close to the minimum detection limit for calling nascent transcription, the odds ratio was close to 1, as expected. As *theta* increased, the odds increased steadily towards a 3-fold boost in nascent RNA intensity (**Figure 5H**). These data show that non-specific, loose clustering of enhancers is significantly correlated with transcription, in terms of both burst frequency and burst size.

## Discussion

Enhancer clustering has been invoked to play important roles in diverse molecular processes, including cell fate specification ^5,7,60^, cancer ^9,60^, emergent behaviors like liquid-liquid phase separation (LLPS) ^4,8,55^, and more ^7,8^. Here, we focused on super-enhancers as a strongly supported model of enhancer clustering, using a comprehensive list of over three-hundred-and-sixty SEs defined in mouse embryonic stem cells ^10,15–18^. We found that SEs rarely form contact-clusters, but are frequently found in looser, contact-free associations, which we termed “communities”. SE communities form cooperatively, are largely promiscuous, and correlate with genomic proximity to other enhancers, higher levels of Pol2, Mediator, and positioning near speckles. Furthermore, genes downstream of individual SEs exhibit larger more frequent bursts when their SE is in a larger community.

Recent models of enhancer-clustering suggested two alternate views of enhancer-promoter (E-P) communication. In one view, E-P communication has been viewed as a “private line” in which enhancers don’t generally interfere in one-another’s communication with their target promoter. Alternatively, if enhancers often form contact-clusters, E-P regulation may be better described as some form of “group chat” communication. Our data indicate that at least for SEs, the reality lies between these two. None of the SEs we examined formed 3-way contact-clusters in more than 5% of cells indicating a pairwise E-P mode of regulation rather than “group chat”. However, the observation that SEs often form communities, and that community size correlated with transcription prompts a third model, in which communication is more like a “cocktail-party conversation”: Here, communication of a given SE-P pair is largely ‘private line’, but is regularly amplified by nonspecific/promiscuous interactions from the crowded environment, in which no other SE, taken alone, contributes a substantial effect.

Our data show that communities occur at length scales that are beyond the typical enhancer promoter interactions: 400-700 nm rather than 200 nm. Recent work has offered several hypotheses about how such clustering-without-contact might occur in the nucleus, as well as hypotheses about what it might mean for transcription ^4–9,55^. A particularly popular and controversial model hypothesized that enhancer clustering might elevate the local concentration of transcriptional activators that bind these enhancers, like Pol2 or Mediator, above the critical threshold needed for LLPS of these factors, and thus achieve distinctive effects on transcription ^55,61^. However, technical challenges have left these hypotheses untested ^4–9^. By quantifying the differences in cluster size (and size-distribution) achieved by hundreds of different SEs across the genome with multi-modal measurement of transcription, our data provided evidence to test these hypotheses. We were surprised to find that POL2 and MED1 occupancy correlated significantly with cluster size. This observation supports the hypothesis that the innate ability of POL2 and/or MED1 to form large condensates may be catalyzed on clustered SEs or may drive SE clustering. Despite the clear statistical signature, we also note that the contribution of POL2/MED1 to enhancer clustering was a minor effect compared to linear genome organization. Furthermore, our data suggest that nuclear position such as migration towards speckles or away from LADs, may have a similar contribution to clustering as POL2/MED1. Thus, our data suggest that diverse factors combine to explain SE clustering, in which LLPS behaviors of SE-binding proteins are only one component.

Future investigation will be needed to distinguish whether SE communities are an adaptive solution that has emerged through positive selection in evolution, or a detrimental source of transcriptional noise that evolution has been unable to purge. Extending the framework we establish here to other cell types, such as terminally differentiated cells or pathological conditions, may uncover conditions in which clustering plays a more prevalent role than observed in stem cells, with clearer links to function. Indeed, elevated degrees of enhancer clustering have been reported in several gene-fusion pathologies including BRD4-NUT ^62,63^, HoxA-NUP98 ^64,65^, EWS-FLI ^66^, consistent with a picture in which clustering may be detrimental. Alternatively, developing animals may exploit the heterogeneity created by cross-talk in promiscuous communities to contribute to diversity and robustness of cell fate specification. In either case, we envision a pivotal role for high throughput microscopy approaches like we report here in further untangling the complex interactions in the crowded nuclear environment.

## Acknowledgments

We thank Ana Pombo (MDC, Berlin) Lonnie Welch (Ohio University), Tony Oro (Stanford) and Ashby Morrison (Stanford) for insightful discussion and feedback. We thank members of the Boettiger and Wang lab for feedback and critical reading of the text. This work was supported by grants from the NIH (U01 DK127419), NSF (EF2022182) and fellowships from the Arnold and Mabel Beckman Foundation and David and Lucile Packard Foundation to A.N.B; the Walter and Idun Berry Foundation postdoctoral grant to A.H.; the NSF Graduate Research Fellowship Program (NSF DGE-1656518) to D.J.L.

## Author Contributions

D.J.L., A.H., K.C.W. and A.N.B. designed the experiments. D.J.L. and A.H. performed the experiments. D.J.L., A.H., S.G., and A.N.B. performed analysis and interpreted the results. D.J.L., A.H., K.C.W, and A.N.B., wrote the manuscript.

## Supplemental information

Figures S1–S5.

Table S1.

## Methods

### Cell culture

ES-E14TG2a (E14) mouse embryonic stem cells (mESC) were obtained from ATCC (CRL-1821) and adapted to grow without the assistance of feeder cells. E14 cells were cultured in media containing 500 mL of KnockOut DMEM, 75 mL ES-grade FBS, 5 mL of MEM NEAA, 5 mL HEPES (1M), 500 uL 2-Mercaptoethanol (55 mM), 5 mL GlutaMAX, 5 ml of penicillin/streptomycin, and 50 uL of mouse LIF (10^7 U/mL). 6-well plates or 10 cm dishes were coated with 0.1% gelatin for at least 10 minutes at RT before use and washed once with 1X PBS. Cells were passaged every 48h at a ratio of 1:5-1:10.

For all ORCA experiments, cells collected from live culture were detached from plates with trypsin/EDTA and stored in 70% ethanol for up to 3 months ^67^.

### Selecting SEs for Oligopaint probe design

A total of 376 target SE’s were chosen by including all 231 mESC-related SE’s from Whyte *et al*. (2013)^19^ and an additional 145 SE’s, sorted by chromosome, from Sabari *et al* (2018) ^20^. For probe design, a standardized target region of 15 kbp was chosen for every SE, calculated by choosing a region 7.5 kbp 5’ and 7.5 kbp 3’ of the center of the published SE coordinate region (mm9). For SEs with overlapping regions in both publications, the coordinates from Whyte *et al.* (2013) were used. Since the SEs for *Hist1h2bg* and *Hist1h2ad* were within 15 kbp of each other, the SE regions were merged and re-labelled as *Hist1h2bf*. The SEs for *Snord2* and *Eif4a2* were also merged due to linear proximity and labeled under *Eif4a2*. The resulting coordinates were then converted to mm10 using the UCSC Liftover tool, to facilitate comparisons with other datasets mapped to mm10.

### Linking SEs to genes

Gene assignments for each SE were derived from their original publication, Whyte et al. (2013) or Sabari et al. (2018). Briefly, both studies linked genes based on the nearest, expressed RefSeq transcript to the center of each SE. In this study, SEs linked to *Mycn*, *Sox2*, *Tgif1*, *Smad7*, *Kdm6b*, *Spry4*, *Klf9*, and *Nanog* were assigned a hyphenated number to indicate ranked proximity from the center of each SE to the TSS of the main transcript of each gene – 0 for overlap with promoter, and ascending integers > 0 for increasing distance (e.g. Nanog-0 is the SE overlapping with the *Nanog* promoter and *Nanog-1* is the next nearest SE to *Nanog*). Additionally, SEs originally assigned to *Dppa3*, *Slc2a3*, and *Phc1* were relabeled as *Nanog-1*, *Nanog-2*, and *Nanog-3*, respectively, based on published work ^24,44^.

### Oligopaints probe synthesis, sample preparation, and primary probe hybridization

The probe synthesis was performed as described before ^35,68,69^.

Cells that were fixed and then stored frozen in 70% ethanol/PBS solution were plated on poly-(D)-lysine coated glass coverslips. After 3x washes with 1X PBS, cells were permeabilized for 10 minutes with 0.5% Triton-X solution in 1X PBS followed by 2x washes with 1X PBS.

For DNA imaging preparation only, cells were then incubated for exactly 5 minutes in 0.1M HCl solution, washed 3x with 1X PBS washes afterwards, and then incubated with RNAse A solution for 30 minutes and 37□. After prior treatments, cells were then washed 3x in 2X SSC and then incubated for at least 1 hour at RT in a solution of 2X SSC, 50% formamide, 10% dextran sulfate, and 0.1% Tween 20. This solution was completely aspirated from the coverslip and then replaced with 50 ug of DNA primary probe (10 ug/uL) that was mixed with 30 uL of 2X SSC, 50% formamide, 10% dextran sulfate, and 0.1% Tween 20. A small, 18 mm x 18 mm Sigmacote-treated coverslip was placed on top of the probes and cells, and the entire was incubated on a 90□ heat block surface for 3 minutes. Afterward, the sample was placed in a hybridization chamber and incubated at 42□ overnight. After incubation, the sample was incubated in 42□ 2X SSC for 10 minutes before the 18 mm x 18 mm coverslip was removed. This was followed by another 10 minute incubation in 2X SSC at 42□. The samples were washed 1x more in RT 2X SSC and then incubated in a fixation buffer (8% PFA, 2% glutaraldehyde, in 1X PBS) for 30 minutes at RT. A final 3x wash with 2X SSC was performed before the sample was ready for imaging.

For RNA imaging, cells were prepared similarly to DNA imaging with the exclusion of HCl and RNAse A incubation steps, 90□ incubation steps, and fixation post-42□ incubation. _4_ ug of RNA primary probe was used for incubation. After RNA imaging, cells were washed 3x in 1X PBS, re-permeabilized for 10 minutes with 0.5% Triton-X solution in 1X PBS, and followed by 2x washes with 1X PBS. The DNA imaging protocol was then followed.

### ORCA imaging

Samples were imaged using a custom-built microscopy and microfluidics setup that was previously described ^68^. The microscope configuration is documented Micro-Meta App ^70^ and publicly available through the 4DN data portal: https://data.4dnucleome.org/microscope-configurations/84d52890-7e46-4cf8-810c-071ed0673ce8/.

For cell registration and drift correction between rounds of imaging, we used a fiducial signal. Similar to previous ORCA studies, we designed the fiducial to bind to all primary probes in the library. We designed a FISH fiducial signal adapter sequence to bind was hybridized against the 5’ end of the primary probes to increase the overall nuclear punctate signal in the 545 nm laser channel. Secondary oligos conjugated to dyes that fluoresce in 657 nm and 750 nm were used for tag-specific imaging. Chromatic aberrations between the 657 and 750 channels were measured by performing replicate measurements with inverted labels for 4 of the SE pairs, to accurately capture the aberrations at the appropriate focal plane and context of the cells. These data were fit to a 3D 2nd order polynomial used to correct the aberrations.

### Image analysis to identify SE coordinates in 3D

A new image analysis pipeline was adapted from our earlier ORCA approach ^68^ to accommodate the genome-wide nature of our probeset and the resulting challenges in disambiguating alleles. Rather than use a fixed distance from the center of a target locus as a filter for identifying spots and segregating alleles, the revised approach used the entire nucleus as the search area. The major steps are described below. These were implemented in Matlab(™) R2022a, and the functions and recipes for the analysis are included in the github repository associated with our manuscript. https://github.com/BoettigerLab/SEclustering-2024

<u>Drift correction</u>: First, the 647 channel and 750 channel images from each round of hybridization were drift-corrected by aligning a fiducial probe that labeled the entire nucleus using a function that finds the displacement which maximizes the auto-correlation of each image with the first one taken. Then we used the same fiducial marker to segment the individual cells using the cellpose algorithm ^71^. Fine scale drift correction was then computed on a cell-by-cell basis, to allow for cell motion independent of the global coverslip drift.

Spot finding: For each drift-corrected image of each cell in each field of view and each chromatic channel, we selected the brightest two foci that matched the point-spread function (PSF) of our microscope and fit the centroid of these spots to a 3D Gaussian function using a least-squares approach.

<u>Chromatic correction:</u> After all the aligned spots were identified, we used our chromatic correction replicate images to specify all pairs of spots that were imaged first in 647 and then in 750 in different labeling rounds. Points which had no matched partner within 1 um (due to variation in detection or background signal) were rejected, so that all retained points had a unique matched pair in the other channel. We then used these pairs to compute a 3D 2nd-order polynomial transformation that minimized the distance between all pairs. This function provides a measure of the chromatic aberrations in our microscope remaining after the built-in chromatic corrections provided by our Plan Apo correcting lens. This is shown as a vector field in Figure S1, which also shows the measured 3D displacement before and after correction, which improves from a median of 131 nm down to 63 nm (See **Figures S1A-S1C**).

<u>Quantification of residual measurement error and labeling efficiency</u>: We compared the measured 3D positions of SEs recorded in the first few rounds of labeling and imaging, to their 3D positions measured several weeks later, at the end of ∼400 rounds of imaging, to quantify the accumulated uncertainty due accumulated sample degradation, camera noise, labeling background changes, and uncorrected residual drift. This was a little less than 100 nm (precise values and the distribution of errors are shown in **Figure S1D**). We also computed for each SE the frequency with which both alleles were detected in every cell, and found for most SEs to be detected at 90% of the time (see **Figure S1E**).

<u>Allele assignments:</u> After all points have been identified in 3D to sub-diffraction position, we assigned alleles to chromosomes by minimizing the intra-chromosomal distances between all alleles assigned to the same chromosome. This method does not distinguish maternal vs. paternal alleles but does prevent inter-SE distances being inflated by accidentally averaging in trans-distances as well. The resulting 3D localizations with SE assignments and allele assignments are recorded in the core FOF-CT table (see Data Availability).

### RNA quantification

RNA FISH data was processed and analyzed as described previously ^67,68^. Images were flat-field corrected and then processed with maximum intensity projection (Z-axis). We used the Cellpose ^71^ algorithm to segment individual cells. RNA and DNA channels were aligned by maximizing the cross-correlation of the fiducial images, as described previously (see also Code Availability). mRNA foci were identified with a local-maxima search function to identify the brightest pixels within 1 um of the primary SE linked to the chosen gene. This region was then cropped and fit using a Gaussian PSF to identify the nascent RNA intensity (height above background from the Gaussian fit). Alleles with no detectable puncta within 1 um were assigned a burst intensity value of 0.

### *Sox2* SCR distance analysis

Data were downloaded from the 4DN data portal for Huang et al 2021 ^41^, https://data.4dnucleome.org/publications/4f766397-5ead-416e-b891-1c95391dcd77/#expsets-table, and also collected by us as described previously ^35^. The 3D positions of probes corresponding to the start, middle, and end of the *Sox2* control region super-enhancer from the unmodified allele were identified and compared.

### ORCA data analyses

#### Average SEs per cluster

For each cell, we computed the distance from all SEs to all other SEs as a matrix (**Figure 2D**), and thresholded this matrix at 200 nm to determine the single-cell SE contact map. For each SE we then computed the total number of contacts observed. To correct for missed detection events, we normalized the number of contacts by the detection frequency (generally ∼0.9, see **Figure S1E**. We report the average cluster size across all cells, sorted by size (**Figure 2H**). An identical approach with thresholds up to 1000 nm was used for exploring loose clustering.

#### Average SE cluster size per cell

From the matrix of SE cluster size per cell, computed as described for the average SE per cluster, we then averaged over all SEs to find the average SE cluster size within a cell.

#### SE count in 95th-percentile SEs

SE count in 95th-percentile SEs was computed as for the average SE per cluster, but reporting the 95th percentile largest cluster across all cells per SE, rather than the average.

#### Simulation of uniform, random distribution of SEs

We computed *r_ave*, the average radii of the nuclei in our study from the nuclear data segmented by Cellpose (see above). We then uniformly distributed points through-out a sphere of this radius as follows: *theta* = *2*\**pi*\*rand, *phi*=acos(1-2*rand), *r*=*r_ave*\*rand^(⅓) where rand is a random real number in [0,1], drawn 752 times (once for each enhancer) per cell, for a total of 4000 cells. The spherical coordinate vectors were converted into Euclidean basis vectors *x,y,z*, using the standard definitions, *x*=*r*\*sin(*phi*)*sin(*theta*); y=*r*\*sin(*phi*)*cos(*theta*), and *z*=*r*\**cos*(*phi*). From these vectors we computed the distance maps for all 4000 cells using the matlab function *randMap* = squareform(pdist(*xyz*)).

#### Entropy

Entropy per SE was computed as *entropy* = -sum{ *p(e)* log_2_(*p(e)*)}, where the probability distribution of contacts *p(e)* across the set of SEs was computed by dividing the number of times each SE contacted each other SEs by the total number of times those SEs were observed and normalizing this distribution such that sum{*p*(*e*)} = 1. As noted in the text, if *p(e)* is a flat distribution (with equal probability of encountering any SE) this function reaches its maximum value at log_2_(N), where N is the total number of SEs. An SE that always contacted the same other SE, if it contacted any at all, the entropy function goes to 0.

#### Cooperativity

Cooperativity was measured as follows. For all possible SE triplets, A, B, and C, we computed the number of cells in which A and B, B and C, and A and C were each observed within 600 nm, and normalized by the total number of cells in which each of these pairs were both detected. We call this the observed proximity frequency. We then computed the expected A–B-C frequency assuming that A-B and B-C proximity are independent of each other, by computing the product of the A-B and the B-C proximity frequencies. We then computed the observed A-B-C contact frequency by asking how many cells actually had A-B and B-C proximity, compared to the total number of cells in which A, B, and C were observed. We defined the cooperativity as the ratio of the expected to observed.

#### Cobursting

SEs associated with bursts (see above, RNA quantification) within 600 nm of other SEs associated with a burst were considered as a co-bursting community. For every SE, we computed the amplitude of its burst as a function of the size of its co-bursting community. Note, this means, for communities of 3 SEs associated with bursts, each SE from the one community counts as an example of a burst happening among a community of size 3. For Odds Ratio analysis (see below for more details), we compared the “exposure” of having 1 more burst-associated SE within 600 nm (a coburst community >1) with the “condition” of having a burst size larger than *theta*, and varied *theta* across the observed range.

#### Odds Ratio

The Odds Ratio provides a metric of the odds that a given exposure leads to a given condition. For example, the exposure may be “having a cluster size of two or more SEs” and the condition may be “associated with a bursting gene”. We define the following variables: *a*, exposed and has the condition, *b*, exposed and does not have the condition, *c*, not exposed but still has the condition, and *d*, not exposed and does not have the condition. The Odds Ratio is then given by the ratio OR = (*a*\**d*)/(*b*\**c*).

#### Neural network

Neural network training used the `fitcnet()` function in Matlab® 2023a, using the `standardize` flag to balance the data input and a 10 node, single layer architecture. The small scale of the network was selected to minimize the effects of overtraining on our modest data depth. For training network parameters, we partitioned the 752 alleles that we measured into a training set used for determining model parameters and a held-out 25% of the loci as a test set for model validation. As the training algorithm and partitioning of the data are stochastic, we conducted 100 replicate training rounds using different partitions of the test and training set.

### ChIP-seq analysis

The fastq files underwent quality control and trimming through the utilization of Trim Galore (parameters: -q 10). Subsequently, reads were aligned to the mm10 genome via ^72^ (parameters: -p 4 --very-sensitive). Following alignment, the reads were sorted and deduplicated using samtools. Heatmaps are generated using deeptools, utilizing normalized bigwig files as input. Peaks were called using Macs2 ^73^ with a significance threshold set to a q-value of 0.05. Raw reads covering all super-enhancer regions were calculated using samtools for all ChIP-seq data and then normalized for subsequent analysis.

Published datasets used:

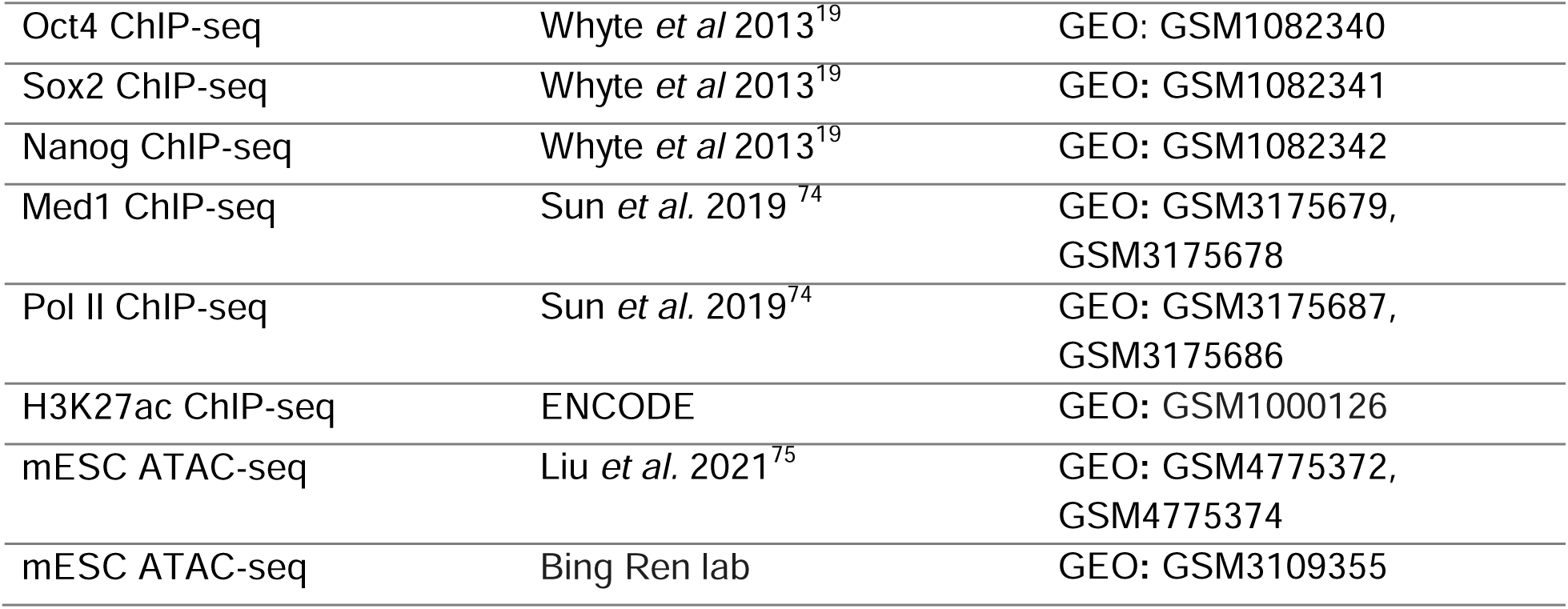

### Processing Hi-C, SPRITE and GAM for ORCA validation

Hi-C data from ^40^ pre-processed contact-read-pairs per chromosome, were downloaded from GEO: GSE161259, and the filtered reads aligning within the 15 kb intervals of our SE regions were extracted for comparison.

GAM data from ^17^ pre-processed per chromosome at 30 kb bin resolution were downloaded <u>GSE64881</u>. The bin with the largest overlap to our SE regions was extracted and used in the comparisons.

SPRITE data from ^18^, pre-processed per chromosome at 20 kb bin resolution, were downloaded from <u>GSE114242</u>. The bin with the largest overlap to our SE regions was extracted and used in the comparisons.

Pearson’s correlation was computed using the built-in Matlab(™) function, corr().

### DamID analysis

DamID data from ^57^, pre-processed to 10 kb bins, was downloaded from GEO, <u>GSE181693</u>. Normalized LAD scores for untreated samples (“PT_0h”) were extracted in 30 kb windows around the bin with the largest overlap to each of our SE regions. Larger windows were included to reduce the effects of data sparsity in the DamID, where resolution is limited by the sequence specificity of the Dam and by the tendency of all SEs to avoid LADs.

### RNA-SPRITE analysis

RNA-DNA SPRITE clusters from ^28^ were downloaded from GEO: GSE186264. The resulting data tables were imported into Matlab for further processing. Clusters containing any of the speckle associated RNAs, Malat1, U1.snRNA, U2.snRNA, U4.snRNA, or U6.snRNA were extracted. For each of these speckle-containing clusters we computed the total number of DNA loci in the cluster. Clusters containing over 100 DNA mapped loci were excluded in order to avoid processing of large fragments containing substantial fractions of nuclei or whole nuclei. The genomic identity of speckle-associated DNAs was mapped genome wide in 250 kb bins, summing all unique reads in each bin.

### Code Availability

Code is available at https://github.com/BoettigerLab/SEclustering-2024

### Data Availability

Data has been submitted for processing at the 4DN data portal. Additional copies of the 4DN formatted data along with several further processed data results are available through Zenodo: DOI: 10.5281/zenodo.10472228. This deposition currently includes:

1. SuperEnhancerLoci.xlsx - a master data table linking the genomic sequence barcode data from the corrected tables (described below) to the corresponding super-enhancer and their genomic coordinates in mm10.
2. Corrected_Data_Tables_by_FOV.zip -- contains drift corrected and chromatically corrected x,y,z coordinates and cellular barcode data to track cell type and coordinate barcode data to identify genomic sequences.
3. Processed_Seq_Data.zip -- re-processed sequencing based data used in this study.
4. FOF-CT_Spot_tables.zip -- draft versions of the 4DN FOF-CT formatted data-standard spot tables. See data format description here: https://fish-omics-format.readthedocs.io/en/latest/
5. Probe_Sequences.zip -- fasta files with the nucleotide sequences of the probes used in the ORCA experiments and multiplexed RNA labeling experiments.

---

**Figure S1.** Quantification of error and uncertainty, related to Figure 1. **A.** Schematic illustration of our approach for measuring chromatic aberrations. **B.** Original and corrected chromatin aberration. Note the correction is more accurate in x,y. **C**. Field portraits of the spatial organization of chromatic aberration. Each graph represents the field of view, each arrow starts at a position of measured chromatic aberration and points in the direction of displacement between the fluorophores. The length of the arrows has been uniformly stretched to be visible on the plot. After correction, the spatially correlated nature of the aberration has been removed. **D.** Quantification of measurement error by replicate labeling. Plot shows the aggregate of all replicate labels, performed up to 376 imaging rounds after the original measurement to capture the greatest extent of potential sample degradation over time. **E.** Detection efficiency per SE barcode. Barcodes are ordered as in Table S1. A total of 14 probes were removed for low detection efficiency and/or off target reactivity and are marked in red.

**Figure S2.** Comparison of ORCA, Hi-C, SPRITE and GAM, related to Figure 1. **A.** Heatmaps of the contact frequency per chromosome for ORCA, Hi-C, SPRITE, and GAM. ChrX is not shown as it contains only 2 SEs. **B.** Correlation of the measured contact frequency between the 4 methods.

**Figure S3.** Effect of community size on entropy and effect of cluster-size threshold on community size, related to. Figure 3**. A.** Correlation of entropy and community size density of SEs indicated by the color map, Pearson’s *r* is shown. **B.** Distribution of average cluster size for each SE, shown as a violin plot, as a function of the cluster radius. Dotted line shows that at cluster-radii greater than 400 nm, most SEs are on average in a cluster, whereas at 200 nm radius all SEs are on average alone. **C**. Correlation plots comparing the effect of cluster radius threshold on the degree of clustering exhibited by each SE. Thresholds of 400-700 all show correlated distributions of cluster sizes with Pearson’s *r* greater than 0.9.

**Figure S4.** Correlates of community size, related to Figure 4. Graphs show the values of the indicated factors which were used to correlate to and predict community size. The x-axes (SE ID) are aligned for comparison between correlates, and are sorted by genome location as in Figure 1.

**Figure S5.** Relation between transcription activity and enhancer communities, related to Figure 5. **A**. Distributions of nascent transcription intensity for Nanog for different community sizes. Pearson’s correlation coefficient and p-values are indicated on the plot and the line over the violins is the best linear regression through all the data. **B**. Number of alleles (number of cells x 2 chromosomes in each cell) for each community size. **C**. Median pairwise distance maps for nascent, transcriptionally ‘on’ and ‘off’ alleles for each of the 6 measured genes.

