## Supplementary figures and images for "Super-enhancer interactomes from single cells link clustering and transcription"

### Supplemental Figure 1

#### Supplemental Figure 1

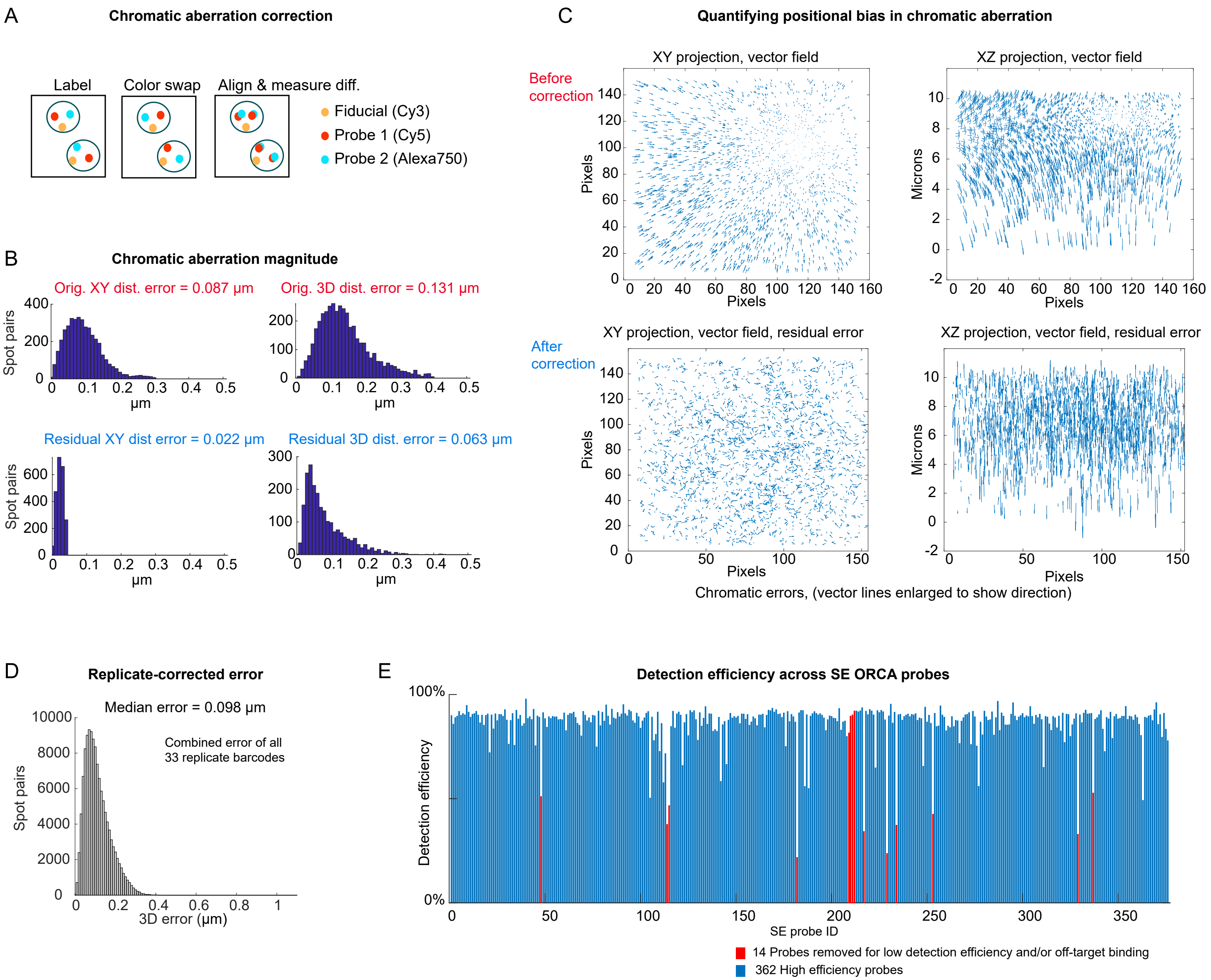

### Supplemental Figure 3

**Supplemental Figure 3**

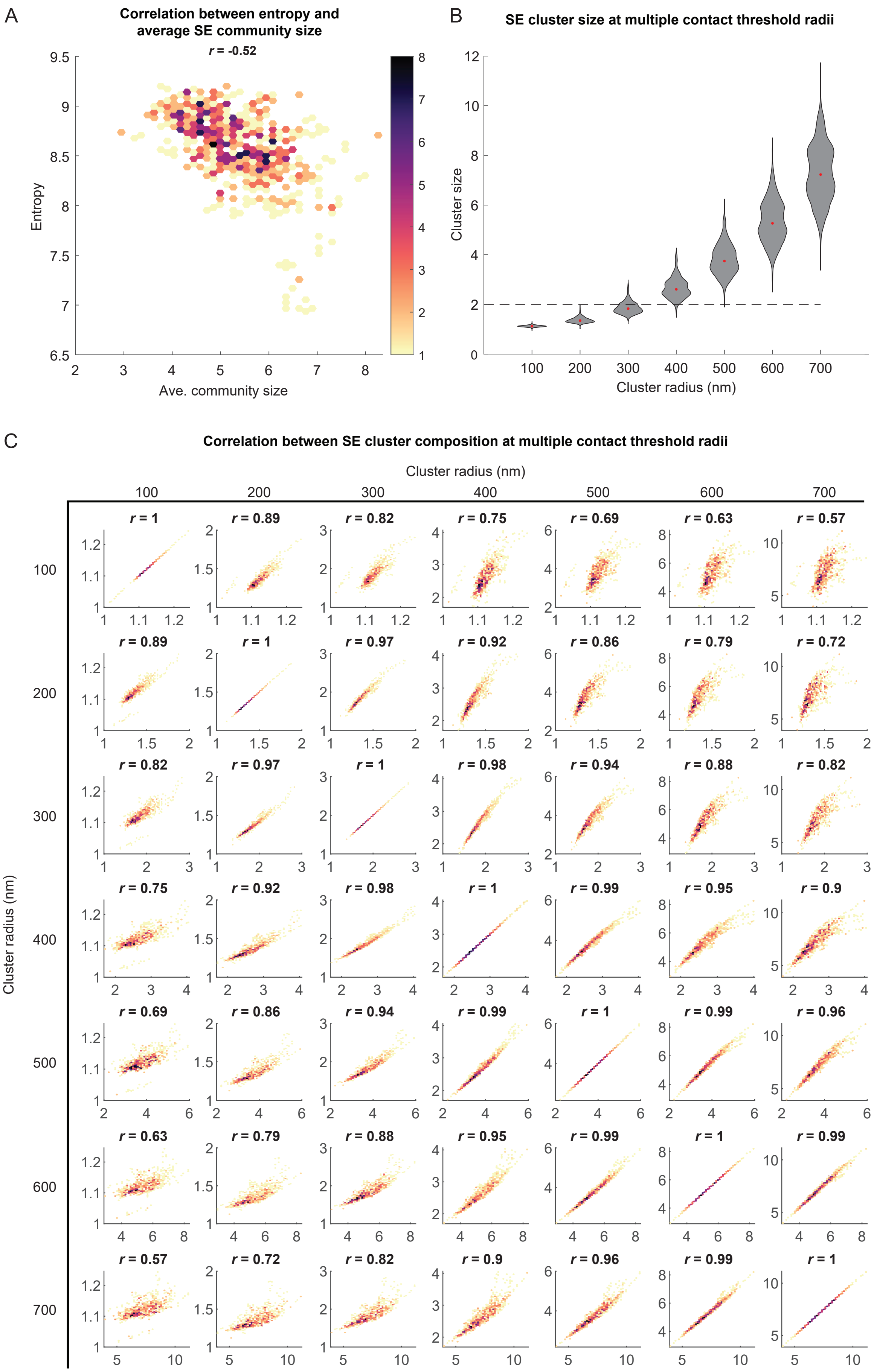

### Supplemental Figure 4

#### Supplemental Figure 4

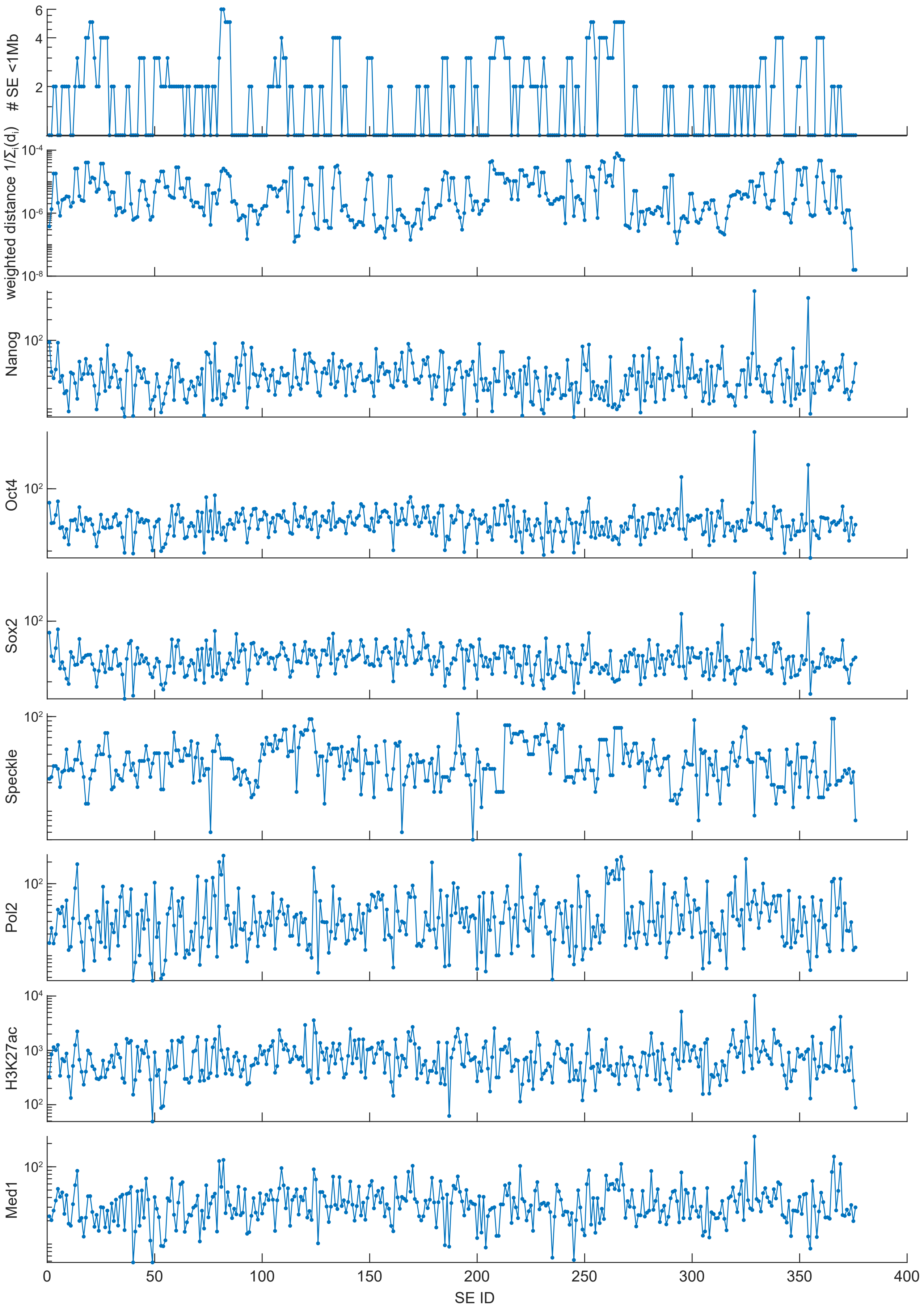

### Supplemental Figure 5

#### Supplemental Figure 5

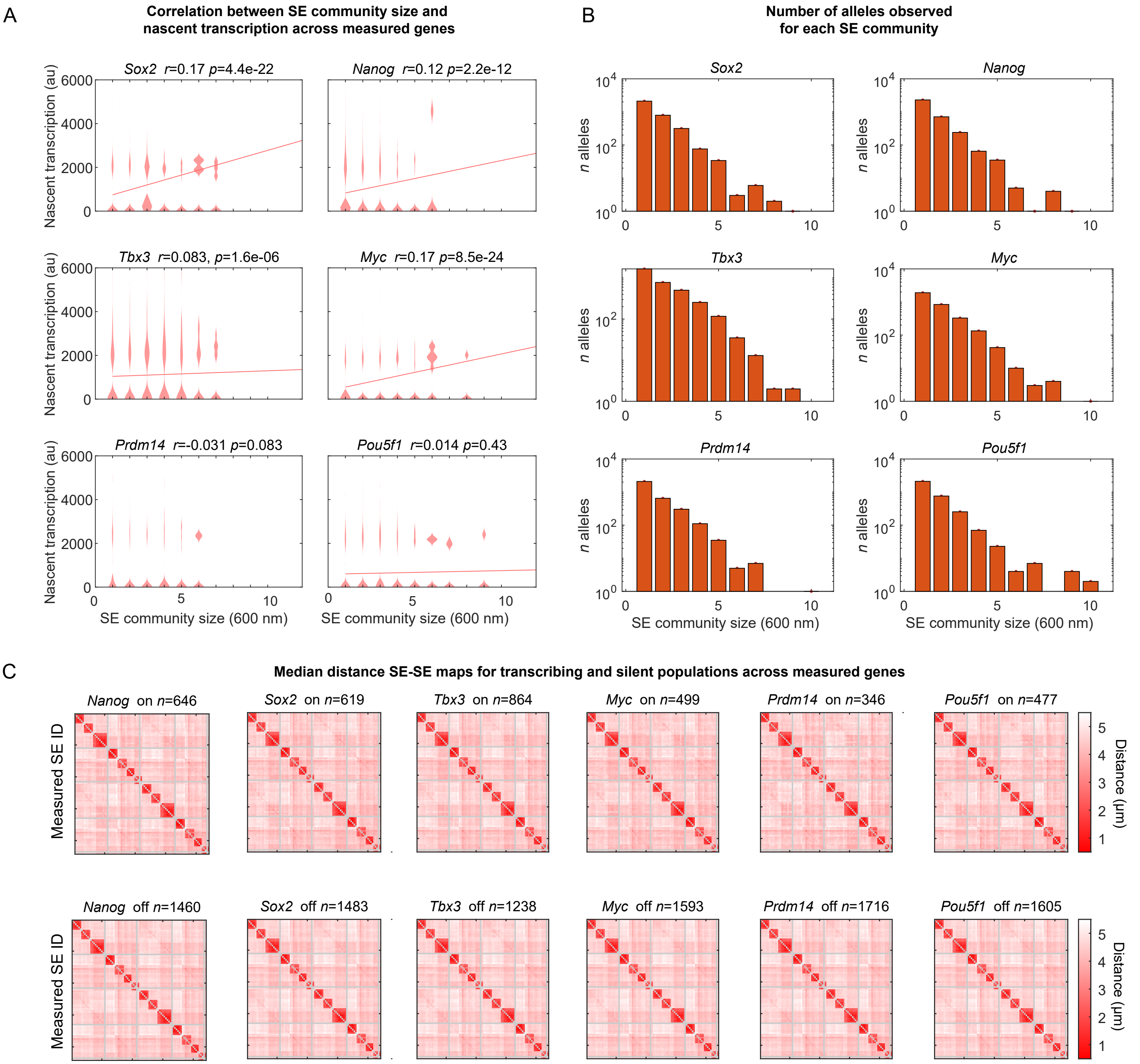
