## Supplemental Figure 2 for "Super-enhancer interactomes from single cells link clustering and transcription"

A

**Heatmap comparison of ORCA with Hi-C, SPRITE and GAM for all chromosomes with >2 SEs**

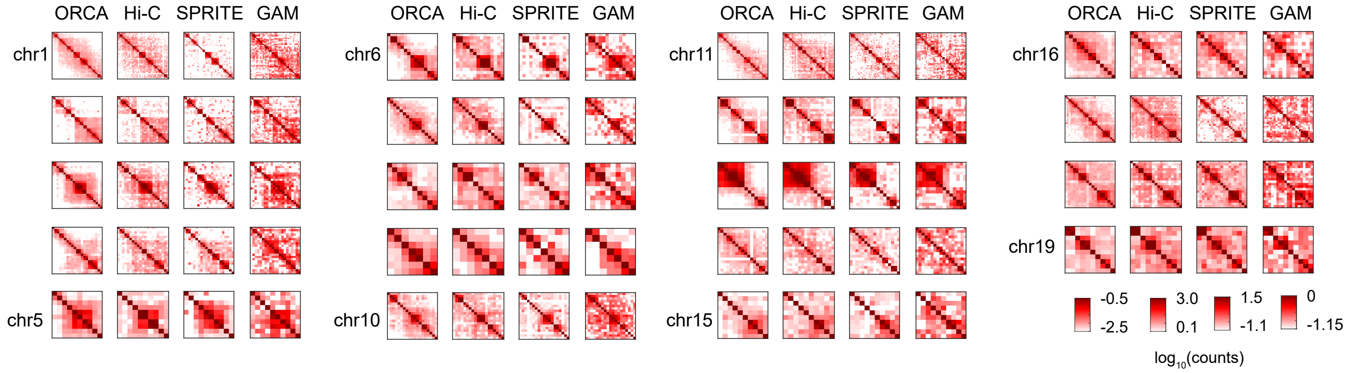


B

**Contact frequency correlations from ORCA with Hi-C, SPRITE and GAM for all SEs genome wide**

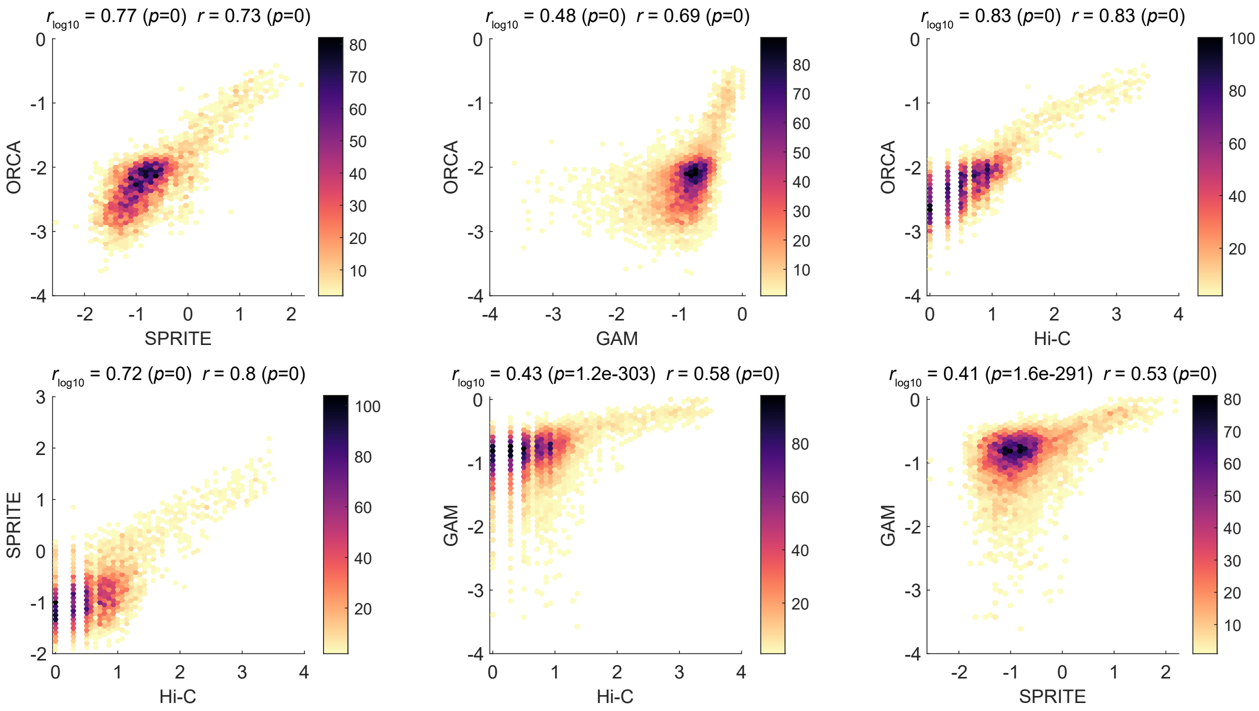
